# Echocardiography-guided percutaneous left ventricular intracavitary injection as a cell delivery approach in infarcted mice

**DOI:** 10.1101/2020.03.31.018358

**Authors:** Yibing Nong, Yiru Guo, Alex Tomlin, Xiaoping Zhu, Marcin Wysoczynski, Qianhong Li, Roberto Bolli

## Abstract

**Background:** In the field of cell therapy for heart disease, a new paradigm of repeated dosing of cells has recently emerged. However, the lack of a repeatable cell delivery method in preclinical studies in rodents is a major obstacle to investigating this paradigm.

**Methods and Results:** We have established and standardized a method of echocardiography-guided percutaneous left ventricular intracavitary injection (echo-guided LV injection) as a cell delivery approach in infarcted mice. Here, we describe the method in detail and address several important issues regarding it. First, by integrating anatomical and echocardiographic considerations, we have established strategies to determine a safe anatomical window for injection in infarcted mice. Second, we summarize our experience with this method (734 injections). The overall survival rate was 91.4%. Previous studies results suggest that 1×10^6^ cells delivered via this method yielded similar retention in the heart at 24 h as 1×10^5^ cells delivered via intracoronary or intramyocardial injection. Lastly, we examined the efficacy of this cell delivery approach. Compared with vehicle treatment, cardiac mesenchymal cells (CMCs) delivered via this method improved cardiac function assessed both echocardiographically and hemodynamically. Furthermore, repeated injections of CMCs via this method yielded greater cardiac function improvement than single dose administration.

**Conclusion:** Echo-guided LV injection is a feasible, reproducible, relatively less invasive and effective delivery method for cell therapy in heart disease. It is an important approach that could move the field of cell therapy forward, especially with regard to repeated cell administrations.

## Introduction

Despite two decades of intensive research in cell therapy for heart disease, the optimal cell type, dosage, delivery method and frequency are still elusive, as is the mechanism behind the observed cardiac benefits of cell therapy[1, 2]. The available evidence supports two fundamental concepts: i) the majority of cell-related effects do not result from direct cardiomyocyte differentiation but rather from paracrine mechanisms[3], ii) all cells, regardless of cell type or delivery approach, fail to engraft in the heart to a significant extent[4]. Therefore, expecting a single administration of short-lasting cells to produce long-term benefits may be unrealistic. A paradigm shift from single administration to repeated administrations of cells is underway[1, 3-5].

Typical cell delivery approaches in clinical studies include intracoronary infusion, transendocardial injection, and direct epicardial injection[1, 2]. The first two are catheter-based, and can be repeated without opening the chest although it would be practically difficult to perform repeated catheter-based deliveries in humans. However, in preclinical studies in rodents, repeated cell administrations can be difficult to perform. Most cell delivery methods used in rodents require open-chest surgery[6-10], and in general, rodents do not tolerate multiple (>2) thoracotomies. Although ultrasound-guided transthoracic intramyocardial injection is a less invasive approach, the efficacy of this method in improving cardiac function remains unclear [11-14]. The uncertainty as to whether the needle tip remains inside the myocardium during the entire injection time (intermittent tip movements would result in cells being delivered into the pericardial space or the LV cavity), the limited targetable myocardial region, and the very limited injectable volume are major challenges that may prevent ultrasound-guided intramyocardial injection from being widely utilized for cell administration.

During our first study of repeated cell administration in mice[15], we established a method for echocardiography-guided, percutaneous injection into the LV cavity as a delivery approach for cell therapy in infarcted mice. More than 700 injections have been conducted in our lab since then, and the method has therefore been standardized. Here, we address several issues regarding this delivery approach. First, we examine the feasibility of the method and the safe cardiac window for injection. Second, we test the reproducibility of the method and discuss how to handle complications. Third, we used previously published data to compare cell engraftment with this method compared with other open-chest approaches. Last, we test whether cells delivered via this method improve cardiac function. We believe this method is an important technical advance that could help to move the field of cell therapy forward, especially with regard to repeated administrations.

## Methods

All animal procedures were performed in accordance with the National Institutes of Health Guide for the Care and Use of Laboratory Animals and were approved by the University of Louisville Institutional Animal Care and Use Committee (protocol number:14034). The general methods have been described previously[15, 16]. Here, we describe in detail the method of echocardiography-guided percutaneous left ventricular intracavitary injection (echo-guided LV injection) and summarize the other methods.

### Ischemia/reperfusion mouse model

Adult, female, C57BL6/J mice were subjected to 60 min of regional myocardial ischemia followed by reperfusion as previously described[15, 17, 18]. In this model, an 8–0 nylon suture was passed under left coronary artery (LCA) approximately 2 mm below the left auricle and a nontraumatic balloon occluder was applied on the artery. Ischemia/reperfusion was induced by tightening and inflating the occluder and then deflating and removing it. To minimize the impact of blood loss, blood from a donor mouse was given at serial times during surgery.

### Determination of safe window for echo-guided LV injection in mice

All echocardiography-related studies were done using a Vevo 2100 Imaging System (VisualSonics, Inc.) equipped with a 30-MHz transducer. Baseline echocardiography was performed on healthy blood donor mice. Parasternal long axis and short axis images were acquired (Fig. 1C and D).

**Fig. 1:**
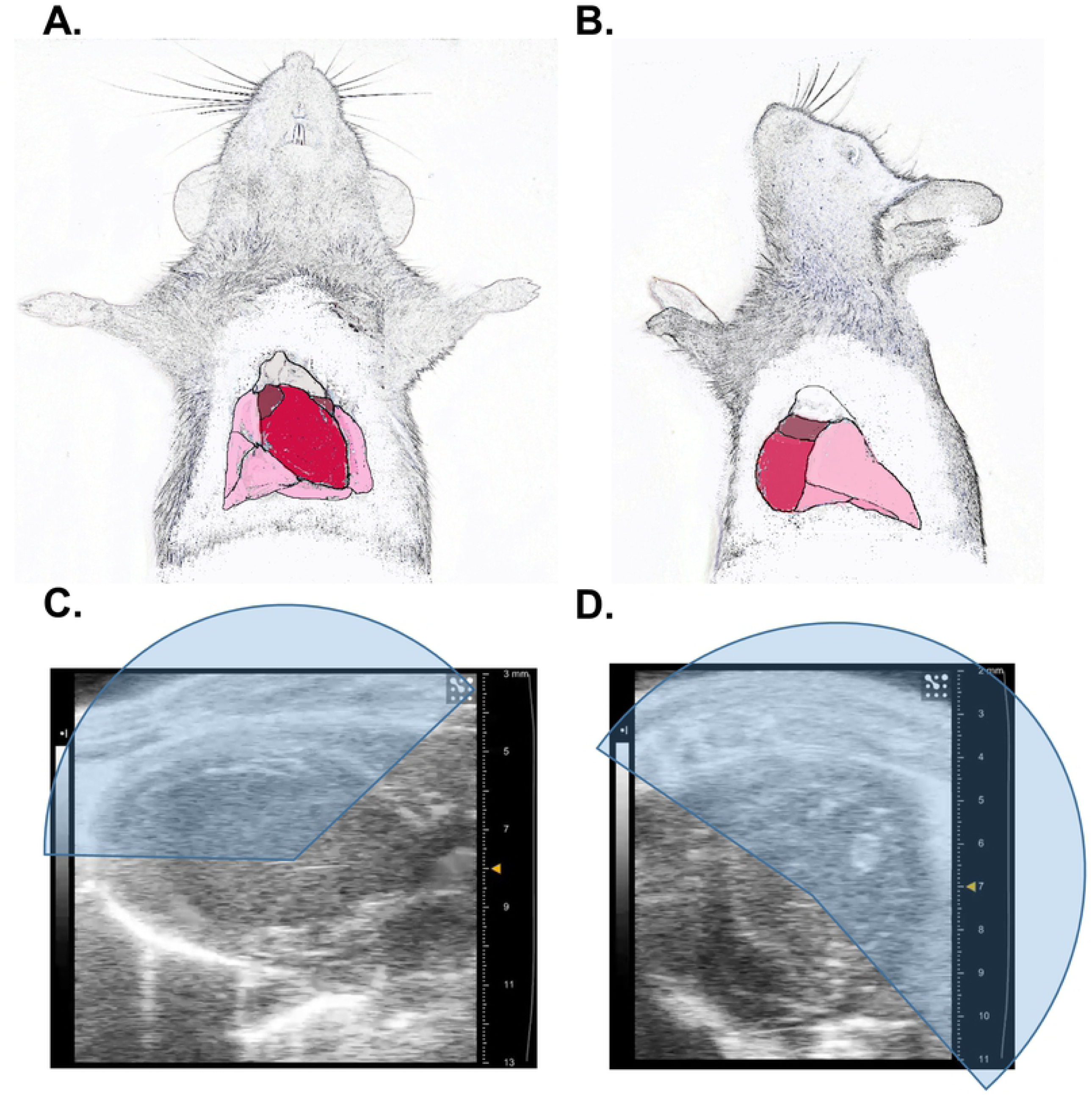
The mouse heart has a wide window for intracavitary injection. Anterior view (**A**) and lateral view (**B**) of the thoracic cage were illustrated in upper panel. Representative parasternal long axis image (**C**) and short axis image (**D**) from a healthy mouse are shown in the lower panels. The fan-shaped shadow indicates a safe window for intracavitary injection without damaging the lung.

After blood was drained via transthoracic cardiac puncture, the ribcage of the donor mouse was immediately removed to expose the heart and major vessels. Heparinized PBS (10 U/ml) was retrogradely infused into thoracic aorta via a 27 G needle attached to a 1-ml syringe. A 5-mm long PE20 polyethylene tubing catheter was inserted into left anterior vena cava and secured by a 5-0 silk suture. The right anterior vena cava and posterior vena cava were then ligated. All three major venae cavae were then disconnected from distal ends. The aorta was then cannulated with a blunted 23 G needle. The heart was sequentially perfused via an aortic cannula with 1 ml saturated KCl, 3 ml heparinized PBS, and 10 ml 4% formalin solution. After fixation, 0.5 ml red latex dye was injected via the aortic cannula and then 0.5 ml blue latex dye was injected via the left anterior vena cava catheter. Digital photos of the heart with visualized coronary artery and cardiac vein were then acquired (Fig. 2A and B).

**Fig. 2:**
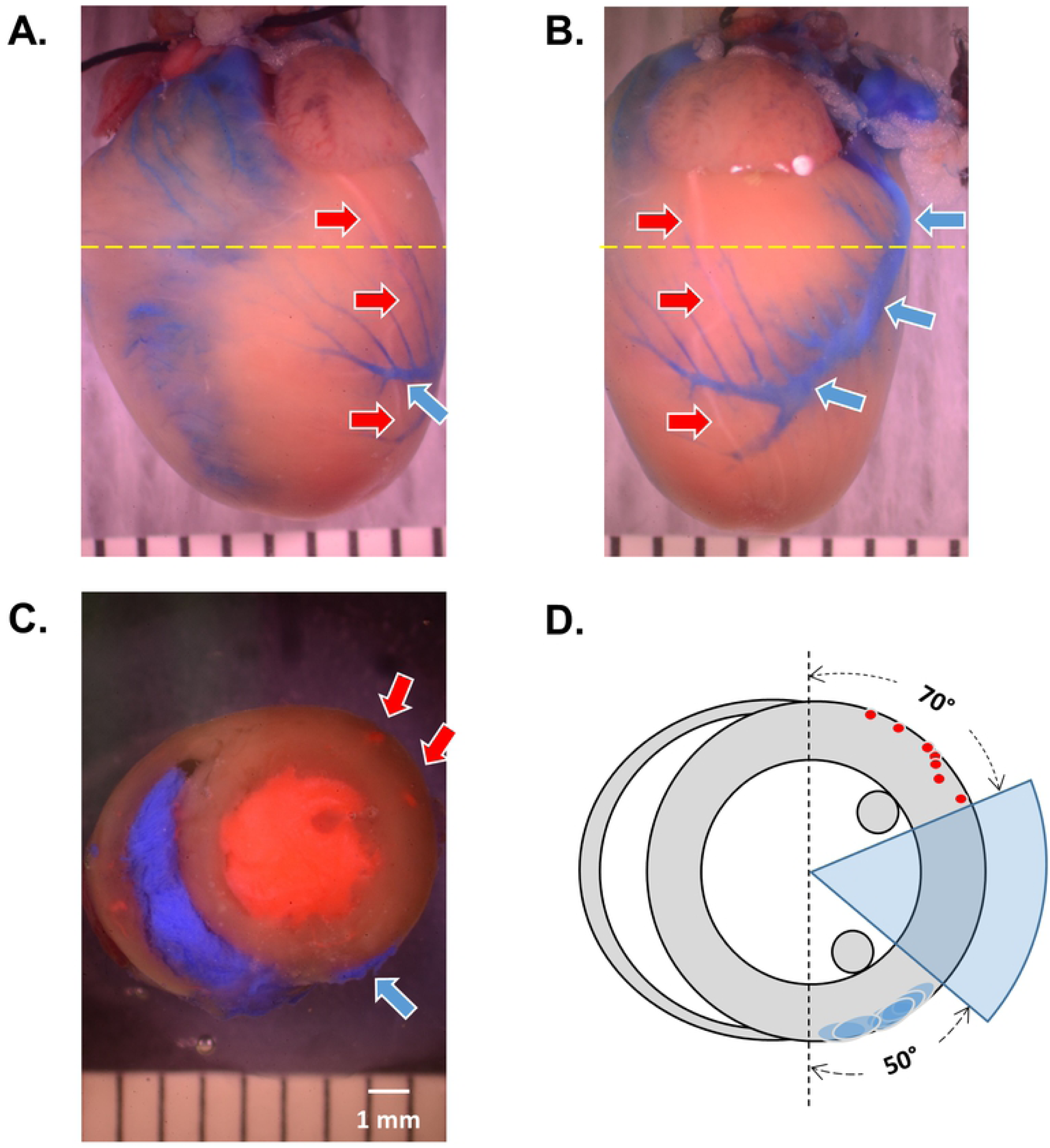
Determination of a safe injection window avoiding major cardiac vessels. Hearts from healthy blood donor mice were fixed and major cardiac vessels were visualized: (**A**) Anterior view of the heart; (**B**) Lateral view of the heart; (**C**) a heart slice was transversely cut at a level 2 mm away from the left auricle (indicated as yellow dashed line in **A** and **B**). Red arrows indicate the locations of left coronary artery, blue arrows indicate the left cardiac vein. (**D**) Locations of LCA (red dots) and LCV (blue ovals) from 6 donor mice. The blue shadow indicates a safe injection window.

The fixed heart was frozen and then cut transversely 2 mm distal to the left auricle. The heart slices were photographed under a microscope. The location of coronary arteries and cardiac veins was visualized in the photo of those slices (Fig. 2C). The photos and the matched short-axis and long-axis echocardiographic images were used to determine the safe window for echo-guided LV injections.

### Echo-guided LV injection in infarcted mice

Successfully infarcted mice that underwent ischemia/reperfusion were subjected to echo-guided LV injection at 3 or 5 weeks after infarction. Some mice underwent multiple injections at 2 weeks or 5 weeks intervals. Cardiac mesenchymal cells (CMCs) or c-kit^+^ cardiac progenitor cells (CPCs) were used in these studies.

#### 1. Preparations before starting

Materials: 30 G 0.5-inch needle, 1 ml syringe, sterile ultrasound transmission gel (Parker Laboratories, Inc.).

Mice were anesthetized with isoflurane (3% for induction and 1.5% for maintenance) and placed on the imaging table in supine position. The anterior chest was shaved and hair removal cream was applied to clean the rest of hair. 10% povidone-iodine and 70% isopropyl alcohol were applied to sterilize the skin. A thick layer of prewarmed sterile ultrasound gel was then applied to the chest wall.

#### 2. Locating the injection window in infarcted mice

The ultrasound transducer was fixed to the mount system in a position such that the transducer was parallel to the forward-backward direction (Y) of the system. The position of the imaging table was adjusted to make sure that a good long-axis view image could be acquired. The transducer was then turned 90° clockwise to the short-axis view position.

Serial short-axis images, 1 mm stepwise from apex to base, were acquired by turning the fine Y direction control knob. In short-axis images, infarcted myocardium could be identified as a thinner wall and with less radial motion or even paradoxical motion (Fig. 3). An experienced echocardiographer can distinguish which level of the short-axis image is above the infarcted zone by carefully reviewing the images. A more accurate but time-consuming method would involve strain analysis to distinguish infarcted and noninfarcted regions (Fig. 4A and B).

**Fig. 3:**
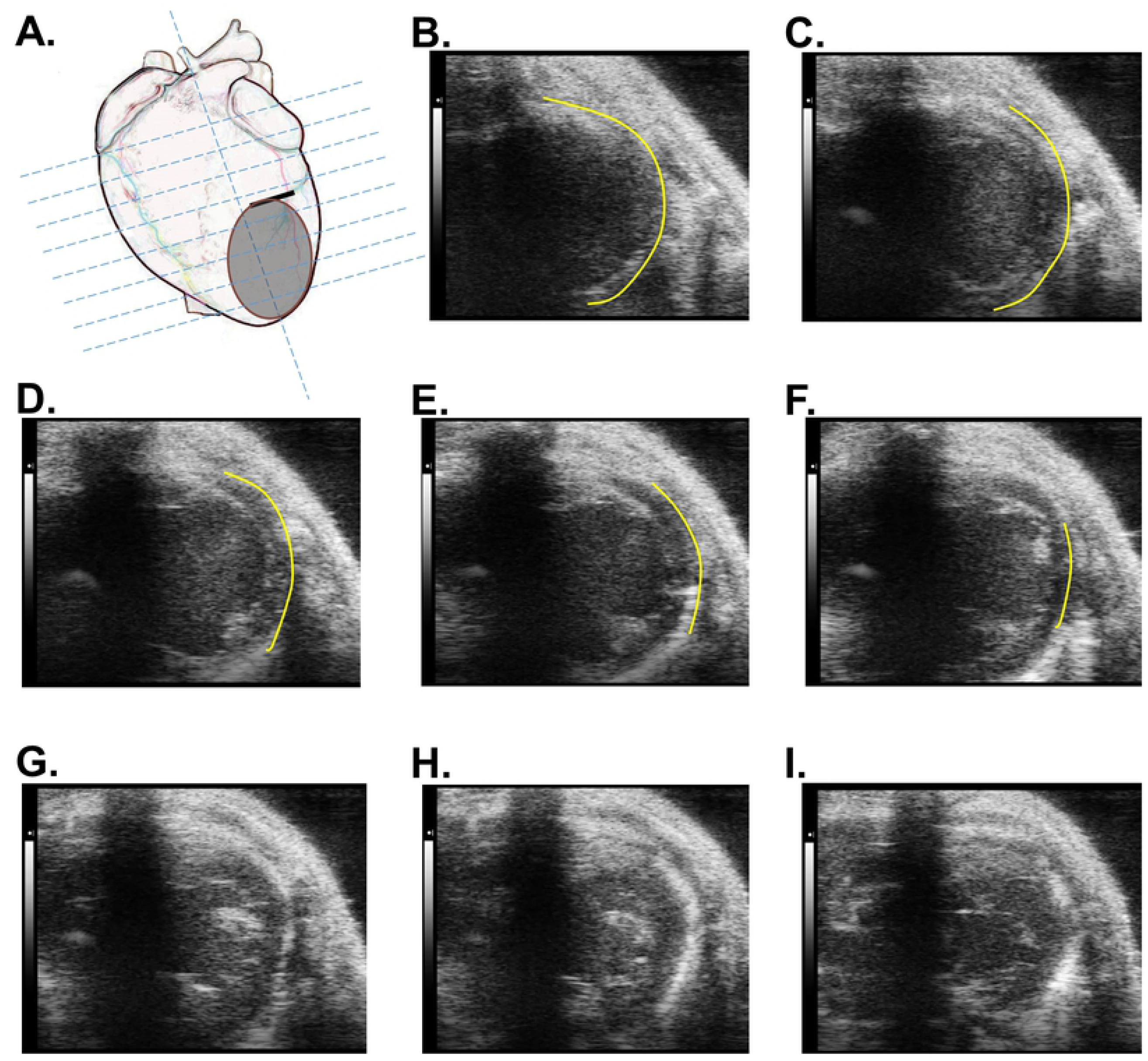
Serial short-axis images from apex to base are shown to determine a safe level for injection. (**A**) Diagram of serial short-axis levels. (**B**) to (**I**) Representative short-axis images, 1 mm stepwise from apex to base, are shown. The yellow line indicates the infarcted zone defined by thinner wall and less radial motion.

**Fig. 4:**
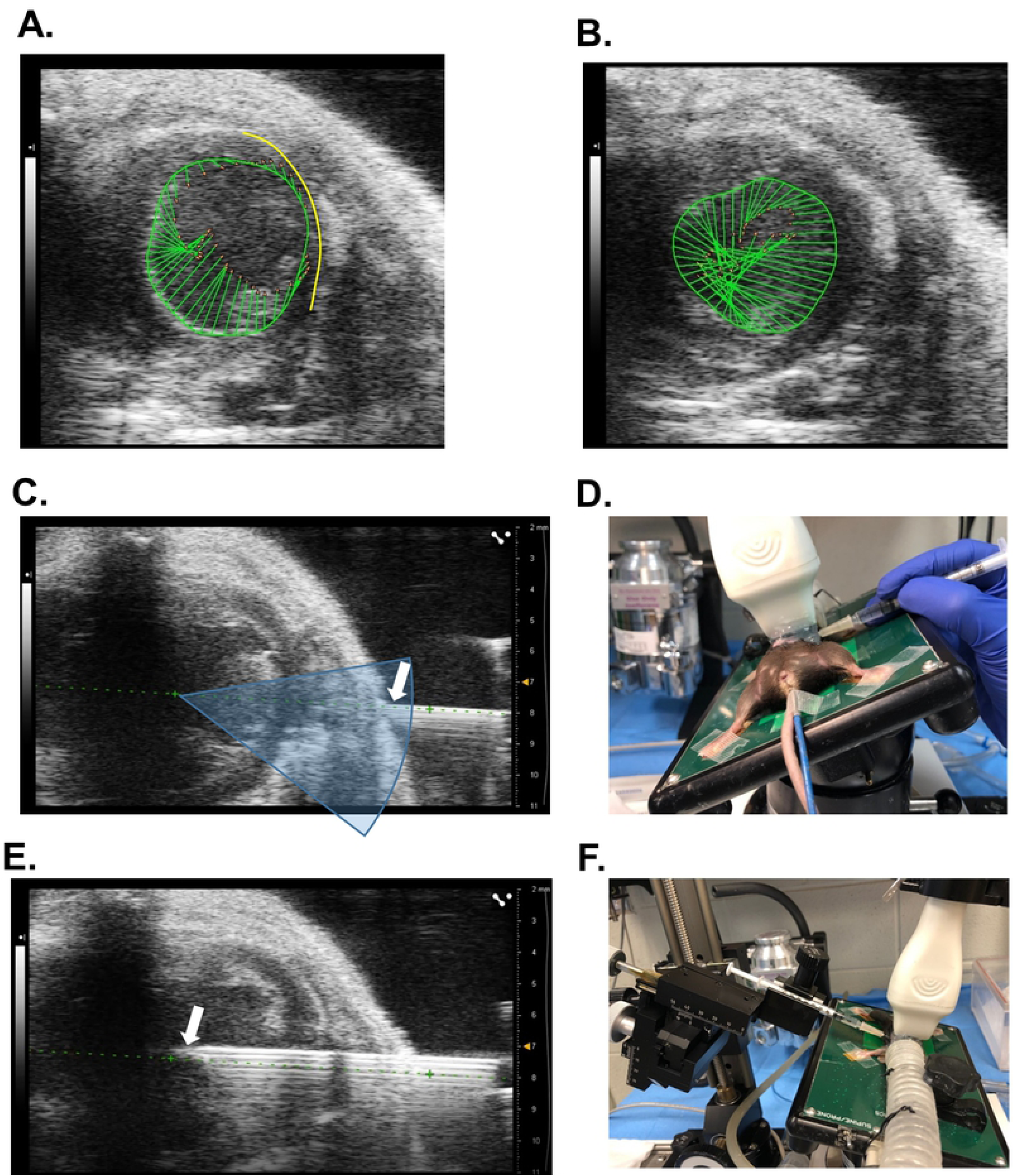
Echo-guided LV injection in infarcted mice. Strain analysis was performed in short-axis images from an infarcted mouse. Vector view at end-systole is shown in the upper panel. (**A**) Vector of radial motion indicates the infarcted zone with reduced or reversed motion. (**B**) Short-axis image above the suture level shows a synchronous myocardial radial motion, indicating a safe level for intracavitary injection. (**C**) Representative echo-guided LV injection image (before insertion). The green dashed line is the guidance line for injection. The white arrow indicates the tip of the needle. The fan-shape shadow indicates a safe window for injection. (**E**) Representative echo-guided LV injection image with injection needle in the cavity. (**D**) and (**F**) Setups for freehand and Vevo Injection System injections.

A short-axis image above the site of the suture placed during ischemia/reperfusion surgery with synchronous myocardial radial motion was considered to be a safe injection level.

#### 3. Injection

Echo-guided LV injection can be conducted by free hand or by using the Vevo Injection system.

For a right-handed operator, the mouse was put on the imaging table with the tail toward the operator. The imaging table was then tilted 30-45° to the right of the animal and the transducer mount was accordingly turned 35-50° counterclockwise to give enough window to access the left lateral chest wall (Fig. 4D).

For a left-handed operator and the Vevo injection system, the imaging table was turned 180° with the head of the animal toward the operator. The table and transducer mount were adjusted accordingly to let the left lateral chest wall face the injection mount (Fig. 4F).

Re-acquired short-axis images were used to locate the safe injection window if the level had been changed during adjustment of the table. Under real-time B-mode imaging, an injection guidance line was drawn to guide the injection (Fig. 4C).

A syringe attached to a 30 G needle was filled with the desired volume of cell solution, making sure there was no air bubble in the system. The syringe was placed under the transducer with the needle exactly parallel to the scan head. The needle was seen in the real-time screen. Following the guidance line, the needle was then advanced into the LV cavity (Fig. 4E and S2 Video).

1×10^6^ cells in 200 µl medium were injected into the LV cavity over 90 seconds. In some mice, a higher dose of 3×10^6^ cells in 600 µl medium were infused over 200 seconds. The same volume of medium served as vehicle control.

The depth of the needle and the physical status of the animal were monitored throughout the injection. After the delivery of cells, the needle was quickly withdrawn. A cotton tipped applicator was pushed against the skin at the insertion site for a few second to prevent bleeding. An experienced operator can finish the whole procedure in 20 min from preparing to completing the delivery.

#### 4. After injection care

Mice were then removed from the imaging table and allowed to recover in a cage on a heating pad. Mice were monitored until able to move freely (typically within 15 min).

### Cell preparation

CMCs[15, 16, 18, 19] and CPCs[7, 8] were isolated and cultured as previously described. All cells were from adult male mice with C57BL/6J background, some were from GFP or RFP transgenic male mice. On the day of injection, cells were harvested, counted and resuspended as previously described. All cells were injected within 15 min after the last centrifugation.

### Echocardiography

Transthoracic echocardiography was performed to assess cardiac function before cell administration and 5 weeks after treatment. In the multiple injections studies, serial echocardiograms were obtained before each injection and at the end of the study. Parasternal long axis images were used to measure cardiac functional parameters[15, 18].

### Hemodynamic study

Mice were subjected to pressure-volume loop assessments right before euthanasia, as previously described[6, 15]. A Millar MPVS ULTRA Pressure-Volume System equipped with a PVR-1035 pressure–volume catheter was used.

### Statistical analysis

Data are presented as mean ± SEM. All data were analyzed with Student’s t tests or one-way ANOVA for normally distributed data followed by unpaired Student’s t tests with the Bonferroni correction, or Kruskal–Wallis one-way analysis of variance on ranks for data that are not normally distributed, as appropriate, A P value <0.05 was considered statistically significant. All statistical analyses were performed using the Sigma Stat software system[20-23].

## Results

### Echo-guided LV injection is feasible in infarcted mice

Unlike the human heart, in the mouse the majority of the LV anterior and lateral wall is not covered by the lung[24, 25]. This anatomical feature gives us a wide window for potential direct transthoracic injection into the LV cavity without damaging the lung (Fig. 1).

To determine a safe window to avoid major cardiac vessels during injections, hearts from 6 mice were fixed and major cardiac vessels were visualized by colored latex dye[26]. On the heart slices cut at 2 mm from the left auricle, the locations of the LCA and left cardiac vein (LCV), including the angles and the distances from epicardium, were measured as illustrated in Fig. 2. Every coronary artery has its unique path and branch tree. But at a distance of 2 mm from the left auricle, the LCA typically has one main trunk[27], and sometimes an early separate branch (1 in 6 observed mice). The distribution of the LCA on the heart slices is also variable but is usually within the LV anterior wall in a 70° arc as illustrated in Fig. 2D. The main trunk of the LCV has less variation and is within a 50° arc in the LV posterior wall (Fig. 2D). Our results suggested that the LV lateral wall between the anterior and posterior papillary muscles is safe to avoid major cardiac vessels.

In infarcted mice, serial short-axis images were acquired to determinate a safe injection level without puncturing the infarct scar. From apex to the suture level, ischemia-damaged myocardium was observed in short-axis images. It manifested thinner wall thickness compared with the remote walls (septal and posterior wall) and either akinesis or dyskinesis. In contrast, above the suture level, about 6 mm from the apex, the non-infarcted myocardium showed a synchronous myocardial radial motion. An experienced echocardiographer can distinguish between infarcted and noninfarcted zone by carefully reviewing the images (Fig. 3). For beginners, strain analysis can be used to determinate a safe injection level[28]. In vector/B-mode view of strain analysis window, the motion vector (direction and speed) of each wall segment is revealed; this made it possible to define the noninfarcted normal motion zone and the infarcted hypokinetic zone (Fig. 4A, B).

The integration of anatomy and echocardiography provided a feasible method for safe echo-guided LV injections in infarcted mice. The method was established during our first repeated cells administration project in mice.

### Echo-guided LV injection is a reproducible, less invasive approach for cell delivery

Since its establishment[15, 16], echo-guided LV injection has been utilized in several projects in our lab. Those projects enrolled 435 mice; among them 181 mice underwent a 3-injections protocol. As shown in Table 1, the overall survival rate after a total of 734 injections was 91.4%. Notably, the survival rate of 181 mice that completed the 3-injection protocol was 81.2%.

**Table 1.** Survival rate after echo-guided LV injection.

|  | Enrolled | Survived<br>after 1<sup>st</sup><br>Injection | Survived<br>after 2<sup>nd</sup><br>Injection | Survived<br>after 3<sup>rd</sup><br>Injection | Total<br>Injected | Survived<br>after<br>Injection |
| --- | --- | --- | --- | --- | --- | --- |
| Single<br>Injection | 254<br>(100%) | 225<br>(88.6%) |  |  | 254<br>(100%) | 225<br>(88.6%) |
| Three<br>Injections | 181<br>(100%) | 151<br>(83.4%) | 148<br>(81.8%) | 147<br>(81.2%) | 480<br>(100%) | 446<br>(92.9%) |
| **TOTAL** | 435<br>(100%) | 376<br>(86.4%) |  |  | 734<br>(100%) | 671<br>(91.4%) |

Autopsies of the 63 mice that died suggested that the major cause of death was bleeding (Table 2). Among 34 bleeding cases, 23 were acute bleeding and the mice died within 5 min after the injections; another 11 mice survived the first 4 h but eventually died 12 h after injections. The majority of the bleeding cases (25 out of 34, 74%) occurred during the first 100 injections, a phenomenon that reflected a “learning curve” for this procedure in our lab. After the method was standardized to avoid major cardiac vessels and infarction scars, only 9 cases of bleeding occurred among the remaining 634 injections.

**Table 2:** Cause of death for all 63 mortality cases.

| Cause of Death |  | Death Cases | Percentage of All Death Cases |
| --- | --- | --- | --- |
| Bleeding |  | 34 | 34/63 (54.0%) |
|  | Acute | 23 | 23/63 (36.5%) |
|  | Chronic | 11 | 11/63 (17.5%) |
| Cell-related |  | 29 | 29/63 (46.0%) |
| Cardiac rupture |  | 0 | 0 (0%) |

Another cause of death was considered to be cell-related because it occurred only in cell-treated mice but not in vehicle groups. Those animals manifested acute respiratory and circulatory failure when the infusion started and most of them died before the total volume of cells could be delivered. Those cases typically occurred at the beginning of a new project. The complications were fed back to the cell preparation lab for quality control. To reduce cell clumping, vials containing cells were inverted several times or gently vortexed before cells were placed into the injection syringe. The injection duration was extended to 90 s instead of the original 60 s. Those changes were sufficient to keep mortality rate under control.

No cardiac perforation or rupture was observed.

In our lab, a new operator with experience in echocardiography has been trained to perform echo-guided LV injections through a training course that required 10-15 mice.

### Cells delivered via echo-guided LV injection improve cardiac function

To determine whether CMCs delivered via echo-guided LV injection are beneficial in chronic heart failure, data from all the projects using the same CMCs and vehicle control were pooled together. Echocardiography assessments were performed 5 weeks after infarction (before treatment) and 35 days after cell administration. Mice were subjected to administration of 1×10^6^ CMCs or vehicle via echo-guided LV injection. There was no difference in LV end-diastolic volume (LVEDV) and EF between the 2 groups before treatment, indicating similar degrees of cardiac dysfunction. While LVEDV did not change after treatment in both groups, CMCs increased stroke volume (SV) and therefore improved ejection fraction (EF) (Fig. 5A-C). Hemodynamic data also showed that compared to vehicle, the CMC-treated group manifested better cardiac function in both load-dependent parameters such as EF and dP/dt (Fig. 5D, E), and load-independent parameter such as end-systolic elastance (Fig. 5F).

**Fig. 5:**
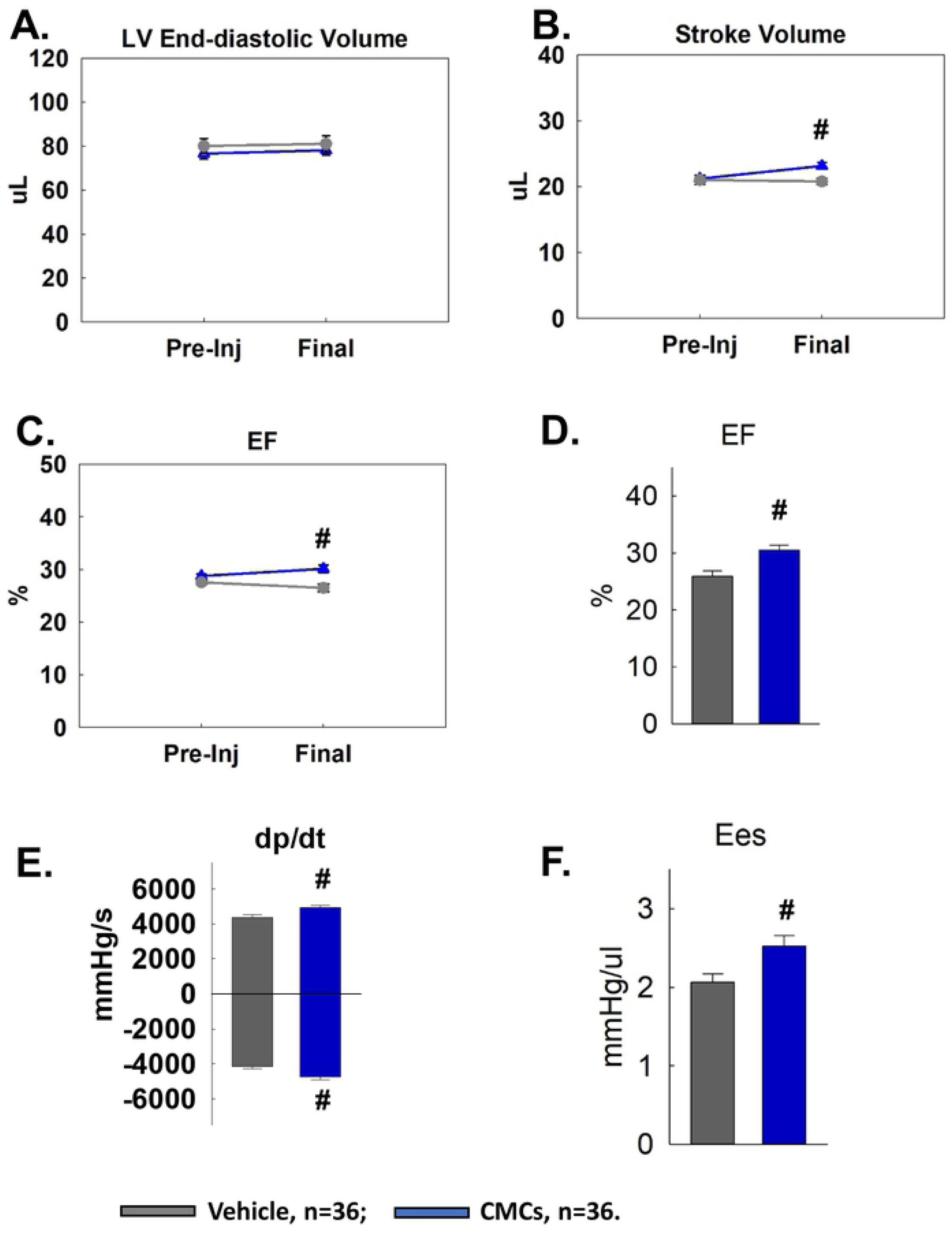
CMCs delivered via echo-guided LV injection improve cardiac function in infarcted mice. Mice with prior MI received CMCs or vehicle, n=36 for each group. Echocardiograms were acquired before and 5 weeks after treatment: (**A**) LV end-diastolic volume; (**B**) stroke volume; and (**C**) EF. Vehicle group in dark grey and CMC group in blue. Hemodynamic assessment was performed before euthanasia: (**D**) EF; (**E**) dP/dt maximum and dP/dt minimum; (**F**) end-systolic elastance. Data are mean ± SEM. **#** p < 0.05 compared with vehicle group.

To test whether repeated administrations of CMCs can produce additional beneficial effects compared with a single administration, we have reported a study in mice with subacute infarction (3 weeks after infarction)[15]. Three repeated doses of 1×10^6^ CMCs were delivered via echo-guided LV injection at 2-week intervals. As reported previously[15], each dose of CMCs resulted in a net increase in EF, such that the cumulative EF gain was greater after multiple doses than after a single dose (Fig. 6B). Therefore, the final EF in the repeated-doses group was significantly higher than in the vehicle or single-dose group[15] (Fig. 6A).

**Fig. 6:**
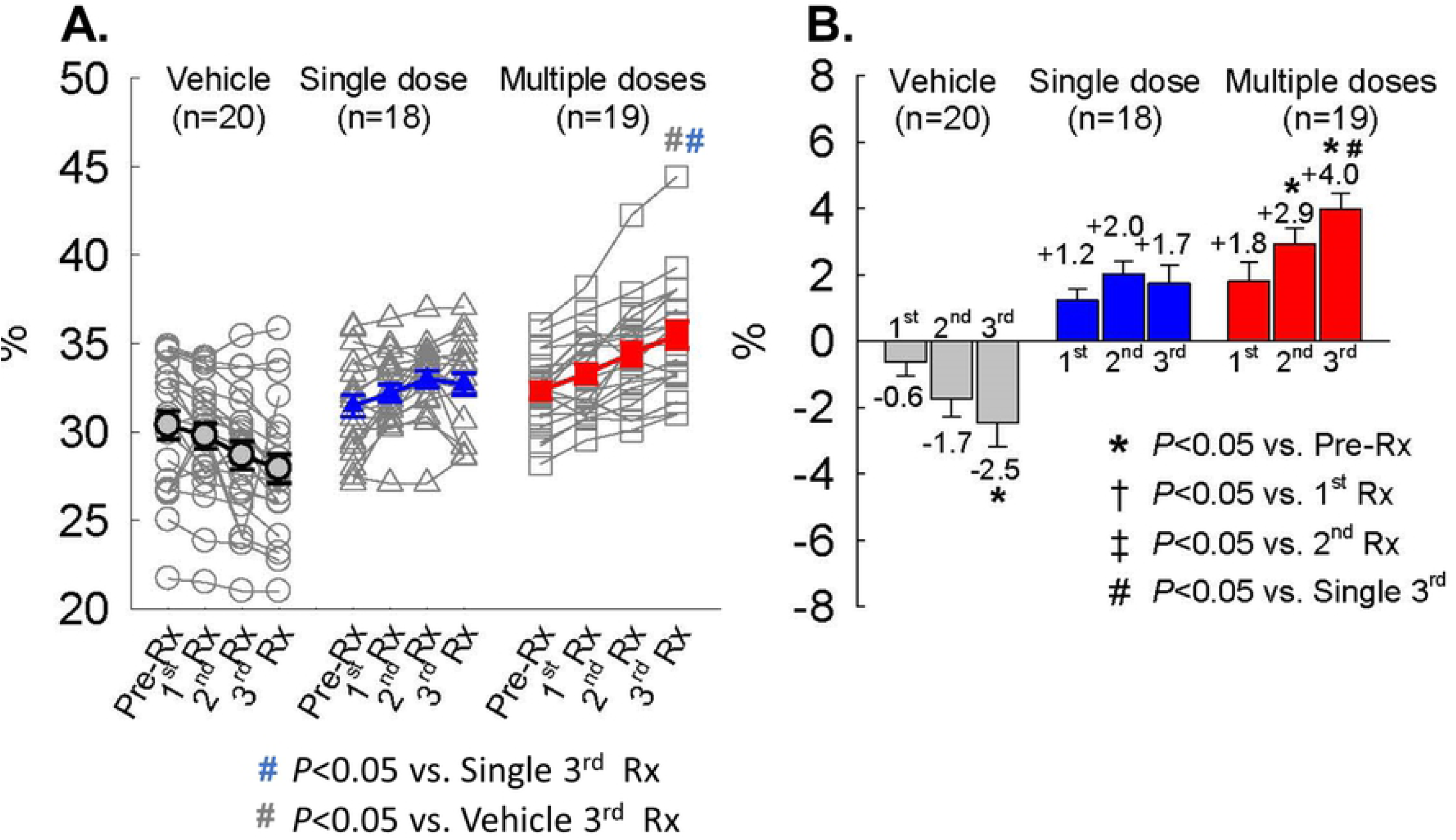
Repeated administration of CMCs via echo-guided LV injection is superior to single administration. EF was measured by echocardiography before treatment (Pre-Rx), 2 weeks after the first treatment (1^st^ Rx), 2 weeks after the 2^nd^ treatment (2^nd^ Rx), and 4 weeks after the 3^rd^ treatment (3^rd^ Rx): (**A**) EF values in individual mice with group mean. (**B**) Cumulative EF gain after each treatment. Data are mean ± SEM. Reproduced from [15] with permission. Copyright ©2017, Springer Nature Switzerland AG.

## Discussion

After two decades of intensive research in the field of cell therapy for heart disease, conclusive demonstration of therapeutic efficacy of the cells is still lacking, and a fundamental understanding of how cell therapy works is still elusive[1, 3, 4, 29-31]. This has led to discussions of changes in the field and search for new paradigms for future research. A paradigm of repeated dosing of cells has become one of the investigational priorities[4, 29, 32]. However, the lack of a reliable, repeatable cell delivery approach in preclinical studies using small animals hinders progress. We have reported briefly on the method of echo-guided LV injection previously[15, 16]. Here, we standardize the method and describe it in detail in an effort to provide sufficient reproducibility for other labs.

Direct intramyocardial and intracoronary injection are commonly used in preclinical studies in rodents. Both of these cell delivery approaches require a thoracotomy, which limits their applicability in studies requiring repeated cells administrations. Some researchers have proposed to use ultrasound-guided intramyocardial[11-14] or intrapericardial injection[33] to deliver therapeutic agents including cells. But in animal models of infarction, the pericardium is not intact and it is uncertain whether the injection needle can be kept in the pericardial space without stabbing the myocardium in a beating heart. Furthermore, pericardial delivery does not allow for cells to infiltrate the myocardium. Echo-guided intramyocardial injections could be repeated in mice. But in an infarcted murine heart, there is limited viable myocardium that could be targeted for injection. Furthermore, because of the thinness of the myocardial wall, the volume that can be delivered is also limited. Finally, it is difficult to be certain that the injection needle remains inside the targeted myocardium during the entire injection without entering the LV cavity. These issues could be the reason why cells delivered via echo-guided intramyocardial injection did not yield cardiac functional improvement[11]. On the other hand, echo-guided LV intracavitary injection allows a larger injectable volume and a larger safe zone without penetrating into the myocardium. The coronary blood flow route also enables cells to reach all cardiac regions. Although this method may require a higher number of cells than direct intramyocardial injection, this disadvantage can be easily overcome. Therefore, we chose echo-guided LV intracavitary injection as a cell delivery approach in infarcted mice.

We first examined the feasibility of this method. Unlike in humans, in rodents the majority of the LV anterior and lateral wall (including the infarcted zone, border zone and remote zone) is not covered by lungs or other organs. By combining anatomical and echocardiographic considerations, we have established strategies to locate a safe window for intracavitary injection and avoid major cardiac vessels and infarct scars. The fact that this method had been successfully utilized in multiple projects in our lab proves its feasibility. A similar method has also been used in a rat model[34, 35] in our group, suggesting its generalizability in preclinical studies of rodents.

Next, we examined the reproducibility of this method. We have used this method in more than 700 injections. As summarized in Table 1, the overall survival rate was 91.4% and the survival rate after 3 injections was 81.2%. The deaths during our “learning curve” were included in the mortality count. With the strategies of locating a safe window and the quality control of cells preparations, bleeding and cell-related complications are expected to be reduced even more in future experiments. Compared to open-chest surgeries that require an experienced surgeon, ventilation, and hours of procedure time, echo-guided LV injection can be conducted by an echocardiographer without using ventilation, and only require about 20 min. In our lab, a new operator with experience in echocardiography can be trained with 10-15 mice. We believe that with the details described herein, it is not difficult for other groups to reproduce this method in their lab.

Third, we discuss here cell retention after echo-guided LV injection. As previously reported[15], to measure cell retention, female mice underwent echo-guided LV injection with 1×10^6^ male CMCs; 5 hearts were harvested at 5 min after injection and another 5 hearts at 24 h after injections. The exact number of male cells in the female heart was calculated as described[36] and compared with our previous studies of 1×10^5^ CPCs delivered via intracoronary or intramyocardial injection[7, 36]. Considering that coronary flow is ∼5% of cardiac output[37] and that cell doses >1×10^6^ may be difficult to produce for repeated injections, we used a 10 times higher dose (1×10^6^ cells) for intracavitary injection to compare with direct intramyocardial injection or intracoronary injection (1×10^5^ cells). The results suggested that intracavitary injection of a 10-fold higher cell dose yielded an almost 2-fold greater cell retention in the heart at 5 min after injection, and similar retention at 24 h compared with the other two approaches (Fig. 7) [15]. Although it is still unclear whether engraftment or retention of transplanted cells is necessary for their cardiac beneficial effects, there is evidence linking the beneficial effects to better retention[38, 39]. Our results may help assuage the concern as to whether sufficient numbers of cells delivered with this method are retained in the heart.

**Fig. 7:**
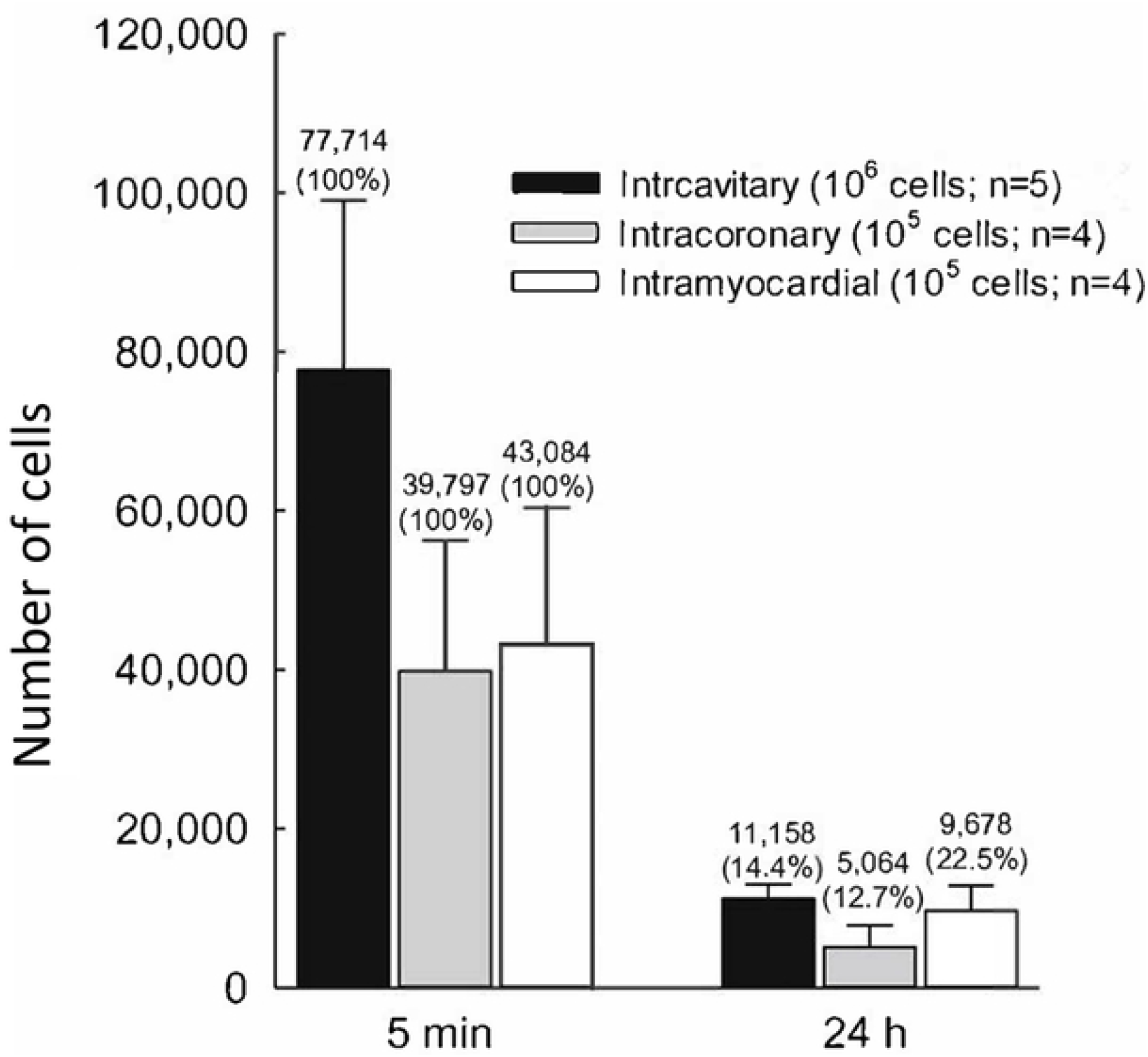
Cells retention in the heart 24 h after administration. Infarcted mice (3 weeks after infarction) were subjected to 1×10^6^ CMC injection via echo-guided LV injection (intracavitary). Hearts were harvested and retained cells numbers were measured at 5 min and 24 h after injection. Data were compared with previous results from intracoronary and intramyocardial injection of 1×10^5^ CPCs at 2 days after myocardial infarction[7, 36]. A highly sensitive and accurate real-time PCR-based method was used to detect the absolute number of male donor cells in a female recipient heart[7, 8, 15, 19]. Data are mean ± SEM. Reproduced from [15] with permission. Copyright ©2017, Springer Nature Switzerland AG.

Finally, the efficacy of this cell delivery approach was examined in the settings of both a single injection and repeated injections. Compared with vehicle treatment, CMCs delivered via this method improved cardiac function assessed both by echocardiography and hemodynamic measurements. Furthermore, repeated injections of CMCs via this method yielded superior cardiac functional improvement than single dose administration[15]. A similar echo-guided LV injection approach was also utilized in rats. As previously reported, repeated administrations of CPCs in rats also produce a cumulative beneficial effect on LV function and structure[35]. Notably, result from rat studies suggested that 3 repeated doses of cells were superior to 1 combined dose even though the total number of cells infused was the same[34]. All these findings support the concept that echo-guided LV injection is an effective cell delivery approach.

In summary, echo-guided LV injection is a feasible, reproducible, relatively noninvasive and effective delivery approach for cell therapy in heart disease. It can be utilized in mice or rat models, at any time point, at any interval, with a wide injection volume capacity and an almost unlimited number of possible repetitions. For example, no other method would enable more than 3 cell administrations in the same animal. It is an important approach that could move cell therapy forward, especially with regard to repeated cell administrations.

## Acknowledgements

None.

## Supporting information

**S1 Dataset. Spreadsheet of cardiac functional data**.

**S2 Video. Representative echo-guided LV injection video**.

